# Predictive distractor processing relies on integrated proactive and reactive attentional mechanisms

**DOI:** 10.1101/2025.04.14.648706

**Authors:** Oscar Ferrante, Ole Jensen, Clayton Hickey

## Abstract

Visual attention is shaped by statistical regularities in the environment, with spatially predictable distractors being proactively suppressed. The neural mechanisms underpinning this suppression remain poorly understood. In this study, we employed magnetoencephalography (MEG) and multivariate decoding analysis to investigate how predicted distractor locations are proactively processed in the human brain. Male and female human participants engaged in a statistical learning visual search task that required them to identify a target stimulus while ignoring a colour-singleton distractor. Critically, the distractor appeared more frequently on one side of the visual field, creating an implicit spatial prediction. Our results revealed that distractor locations were encoded in temporo-occipital brain regions prior to the presentation of the search array, supporting the hypothesis that proactive suppression guides visual attention away from predictable distractors. The neural activity patterns corresponding to this pre-search distractor processing extended to post-search activity during late attentional stages (∼200 ms), suggesting an integrated suppressive mechanism. Notably, this generalization from pre- to post-search phases was absent in the early sensory processing stages (∼100 ms), suggesting that post-search distractor processing is not merely a continuation of sustained proactive processing, but involves re-engagement of the same mechanism at distinct stages. These findings establish a mechanistic link between proactive and reactive processing of predictable distractors, demonstrating both shared and unique contributions to attentional selection.

## Introduction

We are constantly overwhelmed by distractions, from persistent smartphone notifications to frequent advertisement interruptions. Given their prevalence and potential impact on high-intensity goal-directed behaviours – like driving a car or operating heavy machinery – it is essential that we understand how our brain anticipates and minimizes such distractions. In this study, we use multivariate analysis of magnetoencephalography (MEG) to characterize the relationship between proactive and reactive distractor processing when participants ignore distractors that appear at predictable locations.

Visual attention relies on distinct cognitive and neuronal mechanisms for the selection of targets and suppression of distractors (Chang & Egeth, 2019; Van Moorselaar & Slagter, 2020). A particularly striking form of distractor suppression is driven by expectations formed from prior experience (Theeuwes et al., 2022). When salient distractors appear more frequently in a specific location of a visual search array, the attentional priority of that region is reduced, decreasing the interference of future distractors at that location (Ferrante et al., 2018; 2023; Hickey et al., 2014; Sauter et al., 2018; Wang & Theeuwes, 2018). This form of learned suppression occurs without participant awareness of the distractor location manipulation, suggesting a form of statistical learning (Frost et al., 2019; Perruchet & Pacton, 2006). It may reflect a dynamic adjustment of weights in topographic priority maps that guide spatial attention allocation (Britton & Anderson, 2020; Duncan et al., 2023; Ferrante et al., 2018; Ferrante et al., 2023; Liesefeld & Müller, 2021; Wang & Theeuwes, 2018a).

Statistical learning of distractor suppression has been suggested to rely on proactive mechanisms that act before stimulus onset. For example, results show that statistical learning of high-probability distractor locations leads to a reduction in amplitude of the distractor positivity (Pd; Van Moorselaar & Slagter, 2019), an ERP component that appears to index reactive distractor suppression (Hickey et al., 2009). The logic here is that the observed reduction in reactive distractor suppression is made possible by preceding application of proactive suppression. However, until recently, this proactive suppression was not directly identified in brain data, and, as a result, this interpretation has been the subject of debate (Wang et al., 2019).

In this context, we recently tested the hypothesis that statistical learning of distractor suppression reflects a proactive modulation in visual cortex (Ferrante et al., 2023). To do this, we used a recently developed approach to MEG data analysis known as Rapid Invisible Frequency Tagging (RIFT, Seijdel et al., 2023; Zhigalov et al., 2019) to probe neural excitability in visual cortex. Our results showed reduced RIFT response in early visual cortex contralateral to high-probability distractor locations. We concluded that statistical learning of distractor suppression is mediated, at least in part, by the proactive down-regulation of responsivity in early retinotopic visual cortex encoding the high-probability distractor location. Learned distractor suppression thus appears to rely on both proactive (i.e., before stimulus onset) and reactive mechanisms (i.e., in response to distractor salience). However, our understanding of the relationship between these instances of suppression remains rudimentary. Specifically, it is unclear whether proactive and reactive suppression are supported by independent neuronal mechanisms or reflect the same underlying operation that emerges at different times.

Here, we employ multivariate decoding of MEG data to test the hypothesis that proactive and reactive distractor suppression leverage the same underlying mechanism. We first tested whether the location of a predicted distractor could be decoded from pre-search brain activity. We anticipated significant classification of the distractor location in the pre-search time window, reflecting predictive neuronal activity instantiated by proactive attentional mechanisms. We further aimed to characterize the relationship between this proactive mechanism and the reactive processing observed when the distractor is presented in the search display. If proactive and reactive distractor suppression rely on similar attentional mechanisms, we would expect cross-temporal generalization between pre-search and post-search time windows. Conversely, a lack of such generalization would suggest that proactive and reactive suppression are governed by independent mechanisms.

## Materials and Methods

### Participants

The current paper is based on novel analysis of MEG data first reported in Ferrante et al. (2023). Twenty healthy volunteers (13 females) were recruited from the University of Birmingham community. All participants provided informed consent and met local inclusion criteria for MEG studies. The study was approved by the University of Birmingham STEM Ethics Committee.

### Experimental Design

We used a variant of the additional-singleton visual search task (Theeuwes, 1992). Each trial started with a 500-ms fixation point, followed by a placeholder display consisting of four identical non-target stimuli presented equidistantly at 4° of visual angle (v.a.) from central fixation for 1500 ms (Fig. 1A). The non-targets were vertically oriented Gabor patches of size 6° x 6° v.a. displayed in either red or green. After the offset of the placeholder display, a search display was presented for 300 ms. The target stimulus was identified as the Gabor patch tilted 15° to the left or to the right of the vertical meridian. In 66% of the trials, a horizontally oriented colour-singleton distractor (e.g., a red distractor with green target and non-target stimuli) was also presented. Participants had to identify the orientation of the target while ignoring the colour-singleton distractor. A new trial started after an inter-trial interval of between 500 and 1500 ms (random, equal probability distribution).

**Figure 1.**
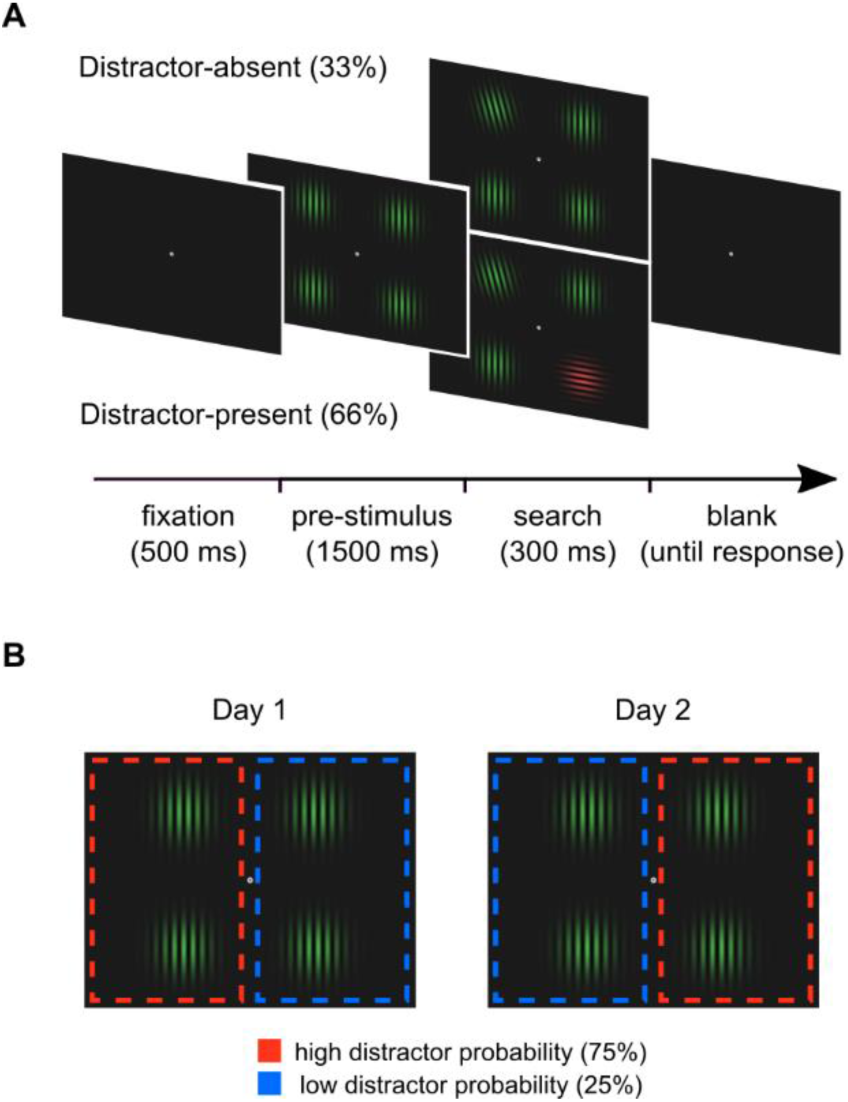
A, Visual search task. Each trial began with a fixation dot, followed by a placeholder display containing four identical Gabor patches. Afterwards, a search display was presented and participants had to report the target orientation (i.e., left or right). In 66% of the trials, a colour-singleton distractor appeared in the search display. B, Statistical learning manipulation. The distractor appeared more often in one hemifield (75%) than the other (25%). Each participant completed two sessions of the experiment (Day 1 and Day 2), with the distractor probabilities swapped between hemifields in the second session (*adapted from Ferrante et al., 2023*).

To induce the statistical learning effect, a probabilistic bias was introduced to the location of the distractor stimuli (Fig. 1B). Specifically, we presented the distractor on one side of the search display in 75% of trials and on the opposite side in 25% of trials. Each participant performed the task in two different sessions, with the distractor probability assignment swapped across sessions. To minimize carryover effects from the first to the second session, sessions were separated by 1-7 days.

As one goal of the experiment was to examine the RIFT signal, as described in Ferrante et al. (2023), frequency tagging was implemented by flickering the two bottom stimuli at very high frequencies (>50 Hz) in the interval from the placeholder display until the offset of the search display. As noted above, these high frequencies are above the flicker fusion threshold, making the tagging invisible. Since our analyses focused on brain activity below 50 Hz, this manipulation is not relevant to the current study.

Participants received verbal instructions at the beginning of each session and completed a practice block consisting of 16 trials. The experiment consisted of 6 blocks, each comprising 120 trials, for a total of 720 trials per session. At the end of the second session, participants filled out a questionnaire to evaluate awareness of the statistical learning manipulation.

### MEG data acquisition and pre-processing

MEG data were acquired using a 306-sensor TRIUX system (MEGIN). MEG data acquisition was conducted following the FLUX Standard Operation Procedure (Ferrante et al., 2022). The data were sampled at 1000 Hz with an online bandpass filter from 0.1 to 300 Hz. Four head position indicators (HPIs) were attached to the participant’s head. The locations of the HPIs, fiducials, as well as the participant’s head shape, were digitized using a Polhemus Fastrack system (Polhemus Inc., USA). Eye movements were monitored using an EyeLink 1000 system (SR Research, Canada). Electrocardiography (ECG) and electrooculography (EOG) signals were recorded along with the MEG data. Manual responses were collected using two MEG-compatible response boxes (NAtA Technologies, Canada).

MEG data were initially converted to BIDS (Niso et al., 2018) using the MNE-BIDS toolbox (Appelhoff et al., 2019) and pre-processed using MNE-Python toolbox (v1.3.1, Gramfort, 2013) in accordance with the FLUX pipeline (Ferrante et al., 2022). Bad sensors were identified and reconstructed using a semi-automatic detection algorithm, after which signal-space separation (SSS, Taulu et al., 2003) was applied to reduce environmental noise. Independent component analysis (ICA) was performed using the *fastica* algorithm (Hyvarinen, 1999). ECG and EOG were used to identify and project out independent components associated to cardiac and ocular artifacts, respectively. Finally, the MEG data were segmented into 4-second epochs, ranging from 1.5 seconds before to 2.5 seconds after the onset of the search display, with time 0 marking the onset of the placeholder display. Epochs with signal amplitudes exceeding an empirical peak-to-peak threshold obtained from the individual MEG data were excluded from the analysis.

### MRI data acquisition and pre-processing

MRI anatomical images were acquired from each participant using a 3-Tesla Siemens Prisma scanner (T1-weighted MPRAGE; TR = 2000ms; TE = 2.01 ms; TI = 880 ms; flip angle = 8°; FOV = 256×256×208 mm; isotropic voxel = 1mm). A single shell boundary element model (BEM) was constructed based on the brain surface derived using FreeSurfer (Dale et al., 1999). The BEM was then used to construct a volumetric forward model (5 mm grid) covering the full brain surface. The lead field matrix was then calculated according to the head position with respect to the MEG sensor array.

### Multivariate decoding analysis

We applied multivariate decoding analysis to the MEG data to classify whether the distractor was presented in the left or right visual hemifield. For classification, we used a linear Support Vector Machine (SVM) classifier from the scikit-learn toolbox (Pedregosa et al., 2011). MEG data were bandpass filtered between 1 and 50 Hz and resampled to 200 Hz. For each time point, the distractor location was classified using data from all 306 MEG sensors within a 25-ms time window (5 samples around each time point, resulting in a feature vector of 306 x 5 = 1530 elements). To further enhance the signal-to-noise ratio, we created pseudo-trials by averaging 5 randomly selected trials without replacement from each of the training and testing sets. To account for the higher number of trials in the high distractor probability condition, we subsampled these trials independently for each participant, randomly selecting a set of trials of the same size as that available in the low-probability distractor location. The resulting data were standardized as z-scores. Classification performance was quantified using the Area Under the Curve (AUC) of the Receiver Operating Characteristic (ROC), which evaluates the classifier ability to distinguish between the two classes (left vs. right). A 5-fold cross-validation approach was employed to ensure robust performance estimates, with the data randomly partitioned into folds. The same procedure was applied for classifying the target location.

To estimate the neuronal sources underpinning each classification activity, we repeated the classification analysis in source space (for a similar approach, see Lu et al., 2024). Source modelling was performed on individual MEG data using depth-weighted minimum-norm estimates (MNE, Hämäläinen & Ilmoniemi, 1994; Wang et al., 1992) combined with dynamic statistical parametric mapping (dSPM, Dale et al., 2000). Epochs were baseline corrected using a 500 ms time window preceding the onset of the placeholder display. Noise covariance matrices were computed over this baseline time window, while covariance matrices for the pre-search time window were computed using data from the onset of the placeholder display to 500 ms after. This was done by first estimating the rank of the data. Then, a common spatial filter was created by combining the baseline and pre-search covariance matrices. MEG data were spatially pre-whitened using the baseline covariance matrix, which allowed for the combination of gradiometer and magnetometer sensors (Engemann & Gramfort, 2015). The MNE-dSPM inverse operator was computed with a loose orientation of 0.2, a depth weighting of 0.8, and applied to the epoched data. The lambda2 regularization parameter was set to 1 (SNR = 1), and the source dipole magnitude was constrained to the normal orientation. Classification was then conducted on the source-level data using the same linear SVM decoder described above. To enhance the signal-to-noise ratio, each time point was classified within a 25-ms time window, data were averaged across every five randomly selected trials without replacement, and the top 500 features were selected through univariate feature selection (i.e., F-test) and submitted to the classifier. The resulting source-level decoding weights were transformed into physiologically interpretable classification patterns (Haufe et al., 2014). Finally, individual source-level classification patterns were morphed to the FreeSurfer averaged brain (*fsaverage*) for group-level comparisons.

We additionally conducted multivariate temporal generalization analysis (King & Dehaene, 2014) to examine the temporal relationship between pre-search and post-search distractor location representations. The temporal generalization relied on multinomial logistic regression with 5-fold cross-validation. The classifier was trained on data from a specific time point and then tested on all other time points across the 306 MEG sensors. Similarly to classification, this analysis was conducted within a 25-ms time window averaged over pseudo-trials generated as mean averages of 5 randomly selected trials.

Cluster-based permutation tests (Maris & Oostenveld, 2007) were conducted to statistically assess above-chance classification performance. Two-tailed one-sample t-tests were applied to each data point using a significance threshold corresponding to α = 0.05. Clusters were then formed by grouping adjacent significant data points. Cluster-level statistics were computed by summing the t-values within each cluster. Statistical significance was calculated using a Monte Carlo null distribution calculated from 1024 permutations of the data with randomization of conditional labels.

### Data and Code Accessibility

MEG data are openly available on request. The experiment code is available at https://github.com/oscfer88/dSL_RIFT/tree/main/experimental_paradigm, and the code for performing all analyses can be found at https://github.com/oscfer88/dSL_RIFT_MNE.

## Results

### Proactive Encoding of Distractor Location in Pre-search Brain Activity

We used temporally resolved classification to determine if the location of distractor stimuli could be identified in the MEG signal. The high-probability location is thought to be proactively suppressed: early visual neurons representing this position show reduced excitability (Ferrante et al., 2023) and when a distractor ultimately appears here, it has less impact on behaviour (Ferrante et al., 2018). We approached analysis with the expectation that this suppression might have a correlate in the pattern of neural activity observed in the pre-search time window, and thus that the distractor location could be identified in classification analysis of MEG data before its appearance.

The results showed significant classification of distractor location in the post-search time window (peak classification time: 205 ms after search display onset; Fig. 2A), indicating distinct brain activity depending on whether the distractor was presented in the left or right hemifield. More importantly, we were able to successfully classify the distractor location during the pre-search time window, with peak classification emerging 175 ms after placeholder display onset (1325 ms before the search array appeared). This suggests that the statistical learning of distractor suppression influenced the proactive encoding of the distractor location, providing predictive information about its likely location in the upcoming search array.

**Figure 2.**
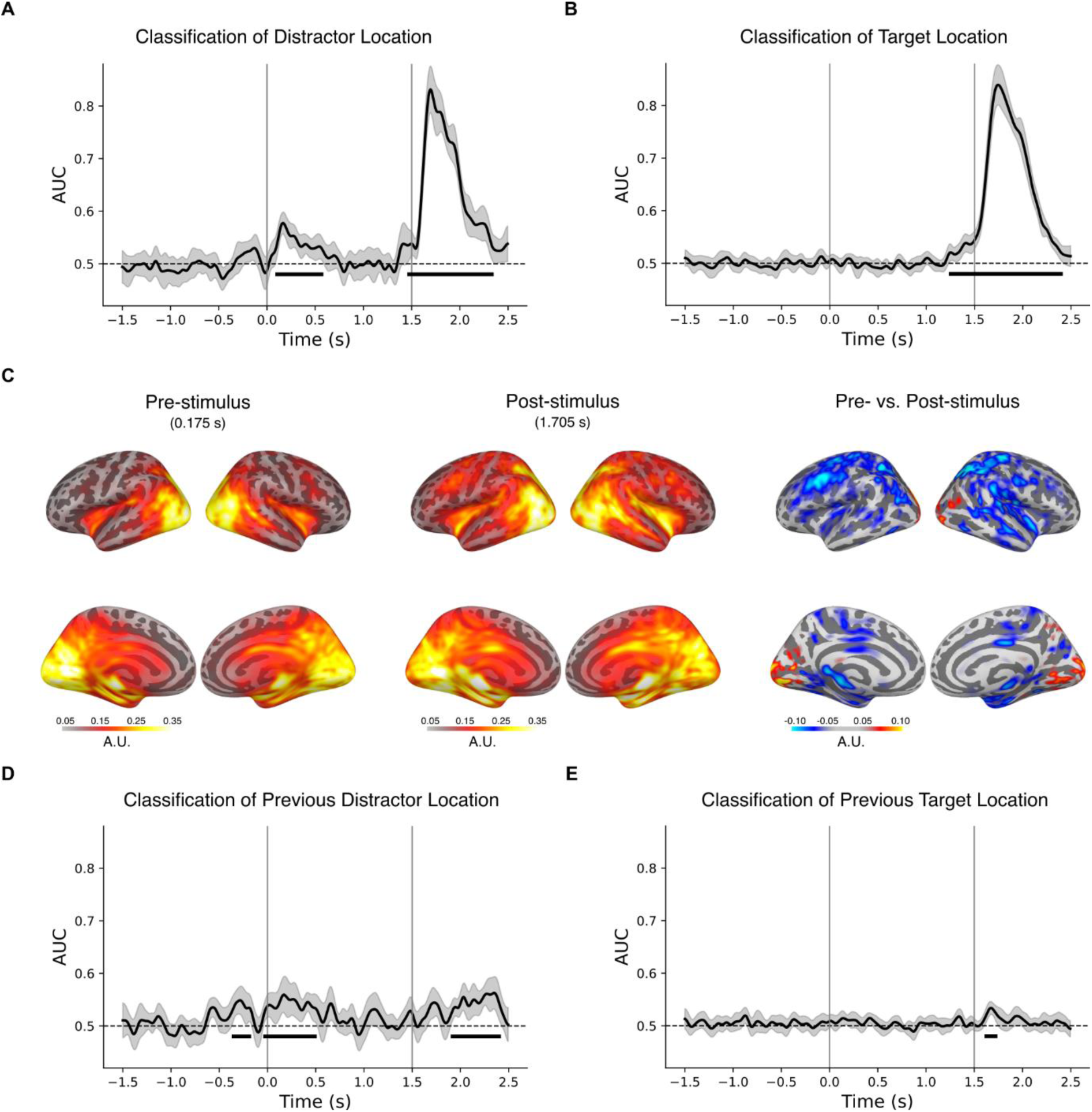
A. Time-resolved classification of distractor location in the current trial (left vs. right hemifield; t = 0 s is onset of pre-search arrays; t = 1.5 s is onset of search array). B. Time-resolved classification of target location in the current trial (left vs. right hemifield). C, Source- space projections of distractor location classification patterns at peak latency for the pre-search period (175 ms), post-search period (1,705 ms), and their difference. Colour bars indicate the level of activation in arbitrary units D, E. Time-resolved classification of the previous (E) distractor and (F) target location. All classification results are presented as Area Under the Curve (AUC) with error bars depicting 95% CI across subjects. Horizontal black lines indicate significant clusters. Vertical grey lines mark the onset of the pre-search (0 s) and the search displays (1.5 s) and dashed black lines represent chance level classification.

The same classification analysis was performed on the target location. Since the target stimulus was presented with equal probability across hemifields, we did not anticipate any proactive representation of its location. In this sense, the classification analysis of the target location serves as a valuable control for the distractor location analysis. For the target location, the classification analysis revealed significant above-chance classification in the post-search time window (Fig. 2B), mirroring the results of the distractor location analysis in the same time window. However, contrary to the distractor location analysis, the classification analysis of the target location did not yield significant above-chance classification in the pre-search time window (Fig. 2B). This distinction was further validated through a cluster-based permutation test comparing the classification time series of the distractor and target locations. Specifically, the pre-search classification for the distractor location was significantly different from that of the target location, with a significant cluster emerging between 95 and 615 ms after the onset of the placeholder display.

Finally, we localized the neuronal sources underlying pre-search distractor processing by replicating the classification analysis on source-level data and computing the corresponding physiologically-plausible classification patterns (Haufe et al., 2014). At the peak of sensor-level classification in the pre-search interval (175 ms), source-level classification identified a bilaterally distributed network across temporo-occipital regions, including the extrastriate cortex, inferior parietal lobule, and middle temporal areas (Fig. 2C, left). We observed additional clusters in medial regions such as the hippocampus and middle-posterior cingulate cortex; however, these findings should be interpreted cautiously due to the challenges of accurately localizing deep sources in MEG. At the peak of sensor-level classification in the post-search interval (205 ms), source-level classification identified a similar temporo-occipital brain network, though with wider distribution, alongside clusters in prefrontal and superior parietal cortex (Fig. 2C, middle). Comparing pre- and post-search source-level classification patterns (Fig. 2C, right), we observed more bilateral activation during the pre-search period, whereas the post-search time window showed greater involvement of prefrontal and superior parietal regions.

### Pre-search Inter-Trials History Effects in Statistical Learning

We further explored how pre-search activity could predict the upcoming distractor location by focusing on inter-trial dynamics. We approached the current data with the idea that inter-trial history may build over repeated experience to instantiate effects associated with statistical learning of distractor suppression.

To test this hypothesis, we trained a sensor-level classifier to identify from data in the current trial (trial n) where the distractor appeared in the immediately preceding trial (trial n-1). Our goal was to determine whether information about the previous distractor location is reinstated in the pre-search activity of the current trial. The results revealed three significant clusters of classification, spanning both pre-search and post-search time windows (Fig. 2D). This indicates that inter-trial history plays a role in both proactive and reactive distractor processing. Notably, pre-search classification was absent when the classification analysis focused on the previous target location (Fig. 2E), suggesting that the pre-search inter-trial effects were not a pure product of repeated experience, but tied to the statistical regularity of distractor location. These results are in line with the hypothesis that statistical learning builds from inter-trial dynamics.

### Proactive and Reactive Distractor Processing Generalize across Attentional Processing Stages

We conducted a temporal generalization analysis to investigate whether the neural features supporting distractor location decoding in the pre-search interval also contributed to its classification in the post-search time window. The confusion matrix illustrating this analysis is presented as Figure 3a. Cluster-based permutation tests conducted on data over the entire time epoch revealed significant clusters in the post-search time window that extended beyond the duration of stimulus presentation (300 ms), indicating distractor processing that continues after the offset of physical stimuli. A focussed analysis of the pre-search time window, from placeholder display onset (0 ms) to search display onset (1500 ms), revealed comparable results during this period. These findings align with the primary classification results above, showing distinct clusters of classification accuracy in both pre- and post-search intervals.

**Figure 3.**
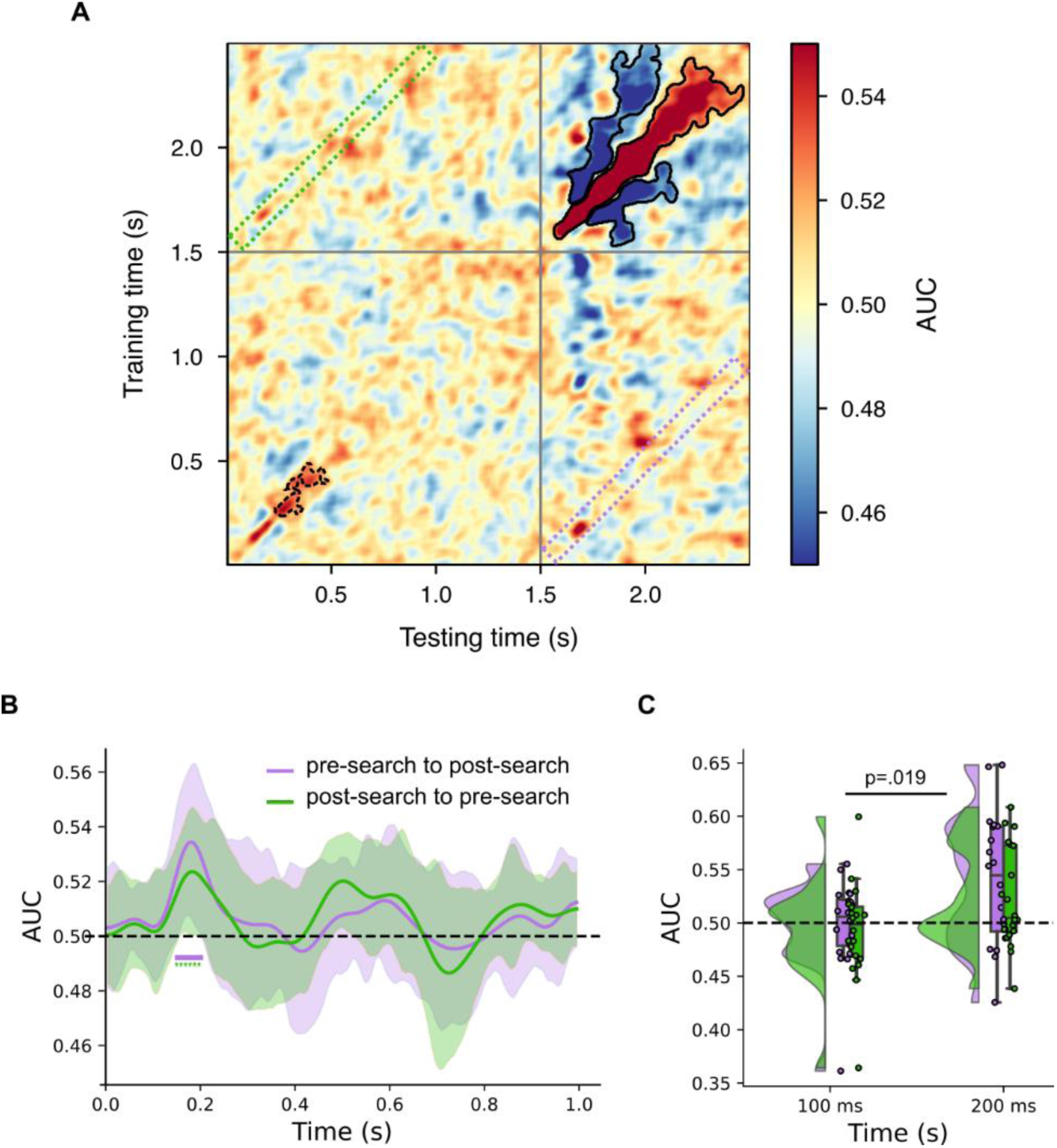
A, Temporal generalization of distractor location classification. Results are presented as Area Under the Curve (AUC). Significant clusters (p<.05) computed over the entire epochs (0-2.5 s) are marked by solid borders. Significant clusters (p<.05) computed over the pre-search time window (0-1.5 s) are marked by dotted borders. B, Cross-temporal generalization between pre-search (0–1 s) and post-search (1.5–2.5 s) time windows. The purple curve represents results obtained by training the decoder on the pre-search time window and testing on the post-search window (illustrated by the purple dotted area in Fig. 3A). For this curve, graph origin reflects onset of a time interval beginning at onset of the placeholder display. The green curve represents results obtained by training the decoder on the post-search window and testing on the pre-search time window (illustrated by the green dotted area in Fig. 3A). For this curve, graph origin reflects onset of a time interval beginning at the onset of the search display. Results are presented as Area Under the Curve (AUC) with error bars depicting 95% CI across subjects. Significant clusters (p<.05) are indicated by horizontal solid lines. Asterisks denote clusters with p<.10. Horizontal dashed line indicates chance level. C, Raincloud plot of cross-temporal generalization centred at 100 and 200 ms delay for pre-search to post-search (purple) and post-search to pre-search (green). Each condition displays the full data distribution, individual datapoints, and the mean with error bars indicating the 95% confidence interval. Horizontal dashed line indicates chance level.

Cross-temporal generalization would emerge in the confusion matrix as off-diagonal clusters of classification accuracy, but no prominent clusters emerge in Figure 3a. However, this broad analysis has low statistical power due to the need for extensive multiple comparison correction. As we approached the experimental data with a priori hypotheses regarding the relationship of pre-search and post-search distractor processing, we conducted additional focussed analyses to further test this idea. Here, we trained a classifier on sequential time windows in the interval following onset of the placeholder display and tested this classifier on corresponding time windows following onset of the search display, and then did the opposite. This meant, for example, that the classifier trained to identify distractor location 200 ms after onset of the placeholder display was used to identify the distractor location 200 ms after onset of the search display, and vice versa. This revealed cross-temporal generalization between pre- and post-search time windows when training on the pre-search and testing on the post-search time window, with a significant cluster emerging with a peak at 185 ms. A very similar result was observed when we trained the classifier on the post-search results and tested it on the pre-search data, with a peak emerging at 175 ms, though this cluster did not independently reach statistical significance (cluster significance: p=0.098).

We conducted a post-hoc analysis on time windows traditionally associated with neuronal correlates of early perceptual (e.g., P1; Clark et al., 1994; Hillyard & Anllo-Vento, 1998) and late attentional processing (e.g., N2pc; Eimer & Kiss, 2007; Hickey et al., 2009; Luck & Hillyard, 1994). Specifically, we performed a series of one-sample t-tests (FDR corrected) on the cross-temporal generalization results, averaging the data over 50-ms time windows centred at 100 ms (i.e., 75-125 ms range) and 200 ms (175-225 ms range), for both directionalities (i.e., training on the pre- and testing on the post-search time window, and vice versa). This identified significant cross-temporal generalization between the pre- and post-search time windows in the 200-ms range (Fig. 3C; pre-search to post-search: t(19)=2.88, p=0.019; post-search to pre-search: t(19)=2.21, p=0.040). In contrast, cross-temporal classification did not differ from chance in the 100-ms range for either analysis direction (pre-search to post-search: t(19)=-0.17, p=0.756, BF10=5.301; post-search to pre-search: t(19)=-0.73, p=0.763, BF10=1.671).

## Discussion

This study examined the neuronal mechanisms underlying the proactive processing of stimulus locations likely to contain distractors. Combining MEG and multivariate analysis, we revealed that pre-search activity in temporo-occipital regions reliably encoded upcoming distractor locations. Notably, this pre-search classification of distractor location generalized to post-search processing in a time window associated with attentional selection (∼200 ms). This suggests that proactive and reactive distractor processing may be governed by shared, integrated attentional mechanisms.

Our findings align with previous research indicating that distractor-specific statistical regularities influence attentional resources (Ferrante et al., 2018; Theeuwes et al., 2022). These attentional effects have been identified in overt behaviour (Ferrante et al., 2018; 2023; Sauter et al., 2018; Wang & Theeuwes, 2018), in modulations of the distractor-elicited Pd ERP component (Hickey et al., 2009; Van Moorselaar & Slagter, 2019; Wang et al., 2019), and in decreased neuronal excitability in visual cortex (Ferrante et al., 2023; Richter et al., 2024; Zhang et al., 2022). Here, we extend this by identifying distinct pre-search and post-search distractor location decoding. Though temporally distinct, our results indicate that these mechanisms share a common neural substrate.

Recent research has explored competing hypotheses regarding how the brain suppresses distracting stimuli (Luck et al., 2021). The signal suppression hypothesis suggests that proactive suppression can occur without initial attentional selection when the distracting features are known in advance (Gaspelin et al., 2015; Gaspelin & Luck, 2018; see also Folk & Remington, 1998). In this context, our pre-search classification results may reflect proactive distractor suppression driven by statistical learning. An alternative hypothesis, known as the “search-and-destroy” model, proposes that distractors cued in advance are initially selected and subsequently suppressed (Klink et al., 2023; Moher & Egeth, 2012; Munneke et al., 2008). This model suggests that improved distractor suppression results from faster disengagement after initial attentional capture (Theeuwes, 2010). In this context, our pre-search classification may reflect the deployment of attention to high-probability distractor locations in preparation for future disengagement. However, this account conflicts with our earlier MEG results (Ferrante et al., 2023), which revealed reduced neural excitability in regions representing high-probability distractor locations. It is difficult to reconcile this reduction in neural excitability with the deployment of attention to these locations, leading us to believe that the current classification reflects distractor suppression that underlies the reduced neural excitability identified in our earlier work.

The pre- and post-search classification results identified above appear to reflect two distinct forms of attentional suppression: proactive suppression, emerging in anticipation of expected distractors, and reactive suppression, emerging as a transient response following distractor onset (Liesefeld et al., 2024; Luck et al., 2021). We approached the current experiment with the idea that these types of distractor suppression might have different physiological correlates, with proactive suppression acting to inhibit early perceptual encoding, and reactive suppression acting to inhibit semantic or task-related representations. This would be broadly in line with results showing that both mechanisms are activated by predictable distractors (Di Bello et al., 2022). However, our results do not provide evidence of this physiological distinction. We found similar patterns of brain activity during proactive and reactive distractor processing, suggesting a shared mechanistic basis.

This impression of shared mechanistic basis was further reinforced by results from temporal generalization analysis. Here, we trained a distractor location classifier on the pre-search time window and tested it on the post-search time window, and vice versa, to find that each classifier accurately predicted the distractor location in both time windows. Notably, this temporal generalization only occurred at a latency around 200 ms after a salient stimulus event – either the onset of the placeholder display in the pre-search interval, or the onset of the search array itself. This is evocative, as neural correlates of attentional engagement and distractor suppression, such as the N2pc (Luck & Hillyard, 1994) and Pd (Hickey et al., 2009), also emerge at this latency. We did not observe temporal generalization at earlier latencies broadly associated with sensory and perceptual processing of visual events (i.e., ∼100 ms from display onset). This suggests that proactive suppression may act by recruiting attentional mechanisms to influence perceptual representation, rather than directly via independent mechanisms of perceptual modulation, at least in the context created by this experiment (Hickey et al., 2023).

We approached the current experiment with the hypothesis that proactive distractor suppression through statistical learning arises from inter-trial dynamics shaped by recent history. When a salient distractor is presented at an unexpected location, reactive suppression reduces its attentional priority to allow attentional redeployment, leaving a transient inhibitory neural trace (Cosman et al., 2018; Ipata et al., 2006). If the distractor reappears at the same location, it elicits a weaker neural response (Grill-Spector et al., 2006) and is less effective at capturing attention (Goschy et al., 2014; Lega et al., 2020; Turatto & Pascucci, 2016). As this is repeated, the inhibitory trace is reinforced and strengthened, ultimately leading to discernible proactive suppression. In line with this idea, our results show that the history of previous distractor locations could be classified from pre-search brain activity in the current trial. Importantly, this pre-search inter-trial effect was not observed for target locations, which were randomly determined. Future research might use similar techniques to track how these intertrial effects build through repetition to generate proactive suppression.

It is worth explicitly noting that pre-search classification of distractor location in our results only emerged after the presentation of the placeholder display. One account for this pattern is that information about future distractor locations is encoded in a silent-activity brain state until it is reactivated by an external stimulus (Stokes, 2015). In this context, the placeholder display used in our task may have acted as a “ping”, reactivating neurons encoding spatial attentional priorities. This interpretation aligns with recent findings from Duncan et al. (2023), who used a manipulation of the target location probability to alter attentional priority (Ferrante et al., 2018; Geng & Behrmann, 2005; Jiang et al., 2013). Through multivariate classification analysis, these authors demonstrated that latent changes in attentional priority could be decoded from neural activity evoked by a “ping” presented during the intertrial interval, suggesting a reactivation of neurons primed by the target spatial regularity. Our findings extend this idea by suggesting that distractor spatial regularities can similarly alter the priority landscape, with this modulation becoming evident when the placeholder display “pings” the representation of distractor locations.

In line with previous literature, we localized the neuronal sources of pre-search distractor location classification to a bilateral temporo-occipital network encompassing occipital, middle temporal, and inferior parietal regions (Hopfinger et al., 2001; Lega et al., 2020; Reeder et al., 2017; Ruff & Driver, 2006; Seidl et al., 2012). Specifically, Ruff and Driver (2006) previously observed increased activity in the middle occipital gyrus in response to a visual cue signalling an upcoming distractor. Recent intracranial EEG evidence from Lin et al. (2024) demonstrates rapid distractor-specific neural processing (∼220 ms post-search) in temporal regions such as superior and middle temporal gyri, amygdala, and anterior cingulate cortex, while revealing weaker contributions from parietal and frontal cortices. Consistent with our current findings, these temporal regions exhibited activity specific to distractor representation, supporting the notion of a temporo-occipital network for distractor processing. Statistical learning of distractor suppression may therefore facilitate the suppression of distractor stimuli by strengthening coupling within the ventral attentional network, a system specialized for detecting salient and unexpected stimuli (Corbetta & Shulman, 2002).

In contrast, post-search distractor classification was associated with prefrontal and superior parietal regions, including the frontal eye fields (FEF) and intraparietal sulcus (IPS). There are at least three possible explanations for this difference. First, proactive suppression through statistical learning may not engage the strategic control mechanisms typically associated with FEF and IPS (Cosman et al., 2018; Ipata et al., 2006; Lega et al., 2019; Marini et al., 2016; Suzuki & Gottlieb, 2013). Instead, it may influence spatial attention through changes in priority maps localized in posterior cortex (Li, 2002; Sprague & Serences, 2013). Second is the possibility that proactive suppression is strategic but relies on a brain network that does not involve FEF and IPS. A final alternative is that the observed proactive effects may be mediated by regions other than FEF and IPS, like the superior colliculus, pulvinar (Krauzlis et al., 2013) and caudate nucleus (Hikosaka et al., 2000), that do not elicit robust MEG signal.

In conclusion, our study demonstrates that predictable distractor locations can be decoded from pre-search neuronal activity in temporo-occipital areas. This proactive representation shares similar mechanisms with the reactive processes triggered at the presentation of the distractor, as evidenced by cross-temporal generalization between pre- and post-search decoding. Furthermore, this temporal generalization is exclusive to late processing stages corresponding to neuronal correlates of attention. Our findings suggest that statistical learning of distractor suppression predictively represents distractor locations, enabling proactive and reactive processing through integrated attentional mechanisms.

## Acknowledgments

OF and CH are supported by the European Research Council under the European Union Horizon 2020 Research and Innovation Program (Grant Agreement 804360 to CH). OJ was supported by a Wellcome Trust Discovery Award (grant number 227420) and the NIHR Oxford Health Biomedical Research Centre (NIHR203316). Our thanks to Jonathan L. Winter and Alexander Zhigalov for providing help with the MEG recordings and to the University of Birmingham’s BlueBEAR high performance computing service (www.birmingham.ac.uk/bear) for computational resources.

## Notes

### Competing Interest Statement

The authors have declared no competing interest.

## References

Appelhoff, S., Sanderson, M., Brooks, T., Van Vliet, M., Quentin, R., Holdgraf, C., Chaumon, M., Mikulan, E., Tavabi, K., Höchenberger, R., Welke, D., Brunner, C., Rockhill, A., Larson, E., Gramfort, A., & Jas, M. (2019). MNE-BIDS: Organizing electrophysiological data into the BIDS format and facilitating their analysis. Journal of Open Source Software, 4(44), 1896. 10.21105/joss.01896

Britton, M. K., & Anderson, B. A. (2020). Specificity and persistence of statistical learning in distractor suppression. Journal of Experimental Psychology: Human Perception and Performance, 46(3), 324–334. 10.1037/xhp0000718

Chang, S., & Egeth, H. E. (2019). Enhancement and Suppression Flexibly Guide Attention. Psychological Science, 30(12), 1724–1732. 10.1177/0956797619878813

Clark, V. P., Fan, S., & Hillyard, S. A. (1994). Identification of early visual evoked potential generators by retinotopic and topographic analyses. Human Brain Mapping, 2(3), 170–187. 10.1002/hbm.460020306

Corbetta, M., & Shulman, G. L. (2002). Control of goal-directed and stimulus-driven attention in the brain. Nature Reviews Neuroscience, 3(3), 201–215. 10.1038/nrn755

Cosman, J. D., Lowe, K. A., Zinke, W., Woodman, G. F., & Schall, J. D. (2018). Prefrontal Control of Visual Distraction. Current Biology, 28(3), 414–420.e3. 10.1016/j.cub.2017.12.023

Dale, A. M., Fischl, B., & Sereno, M. I. (1999). Cortical surface-based analysis: I. Segmentation and surface reconstruction. Neuroimage, 9(2), 179–194.

Dale, A. M., Liu, A. K., Fischl, B. R., Buckner, R. L., Belliveau, J. W., Lewine, J. D., & Halgren, E. (2000). Dynamic statistical parametric mapping: Combining fMRI and MEG for high-resolution imaging of cortical activity. Neuron, 26(1), 55–67.

Di Bello, F., Ben Hadj Hassen, S., Astrand, E., & Ben Hamed, S. (2022). Prefrontal Control of Proactive and Reactive Mechanisms of Visual Suppression. Cerebral Cortex, 32(13), 2745–2761. 10.1093/cercor/bhab378

Duncan, D. H., Van Moorselaar, D., & Theeuwes, J. (2023). Pinging the brain to reveal the hidden attentional priority map using encephalography. Nature Communications, 14(1), 4749. 10.1038/s41467-023-40405-8

Eimer, M., & Kiss, M. (2007). Attentional capture by task-irrelevant fearful faces is revealed by the N2pc component. Biological Psychology, 74(1), 108–112. 10.1016/j.biopsycho.2006.06.008

Engemann, D. A., & Gramfort, A. (2015). Automated model selection in covariance estimation and spatial whitening of MEG and EEG signals. NeuroImage, 108, 328–342. 10.1016/j.neuroimage.2014.12.040

Ferrante, O., Chelazzi, L., & Santandrea, E. (2023). Statistical learning of target and distractor spatial probability shape a common attentional priority computation. Cortex, 169, 95–117. 10.1016/j.cortex.2023.08.013

Ferrante, O., Liu, L., Minarik, T., Gorska, U., Ghafari, T., Luo, H., & Jensen, O. (2022). FLUX: A pipeline for MEG analysis. NeuroImage, 253, 119047. 10.1016/j.neuroimage.2022.119047

Ferrante, O., Patacca, A., Di Caro, V., Della Libera, C., Santandrea, E., & Chelazzi, L. (2018). Altering spatial priority maps via statistical learning of target selection and distractor filtering. Cortex, 102, 67–95. 10.1016/j.cortex.2017.09.027

Ferrante, O., Zhigalov, A., Hickey, C., & Jensen, O. (2023). Statistical Learning of Distractor Suppression Downregulates Prestimulus Neural Excitability in Early Visual Cortex. The Journal of Neuroscience, 43(12), 2190–2198. 10.1523/JNEUROSCI.1703-22.2022

Folk, C. L., & Remington, R. (1998). Selectivity in Distraction by Irrelevant Featural Singletons: Evidence for Two Forms of Attentional Capture.

Frost, R., Armstrong, B. C., & Christiansen, M. H. (2019). Statistical learning research: A critical review and possible new directions. Psychological Bulletin, 145(12), 1128–1153. 10.1037/bul0000210

Gaspelin, N., Leonard, C. J., & Luck, S. J. (2015). Direct Evidence for Active Suppression of Salient-but-Irrelevant Sensory Inputs. Psychological Science, 26(11), 1740–1750. 10.1177/0956797615597913

Gaspelin, N., & Luck, S. J. (2018). The Role of Inhibition in Avoiding Distraction by Salient Stimuli. Trends in Cognitive Sciences, 22(1), 79–92. 10.1016/j.tics.2017.11.001

Geng, J. J., & Behrmann, M. (2005). Spatial probability as an attentional cue in visual search. Perception & Psychophysics, 67(7), 1252–1268. 10.3758/BF03193557

Goschy, H., Bakos, S., MÃ¼ller, H. J., & Zehetleitner, M. (2014). Probability cueing of distractor locations: Both intertrial facilitation and statistical learning mediate interference reduction. Frontiers in Psychology, 5. 10.3389/fpsyg.2014.01195

Gramfort, A. (2013). MEG and EEG data analysis with MNE-Python. Frontiers in Neuroscience, 7. 10.3389/fnins.2013.00267

Grill-Spector, K., Henson, R., & Martin, A. (2006). Repetition and the brain: Neural models of stimulus-specific effects. Trends in Cognitive Sciences, 10(1), 14–23. 10.1016/j.tics.2005.11.006

Hämäläinen, M. S., & Ilmoniemi, R. J. (1994). Interpreting magnetic fields of the brain: Minimum norm estimates. Medical & Biological Engineering & Computing, 32, 35–42.

Haufe, S., Meinecke, F., Görgen, K., Dähne, S., Haynes, J.-D., Blankertz, B., & Bießmann, F. (2014). On the interpretation of weight vectors of linear models in multivariate neuroimaging. NeuroImage, 87, 96–110. 10.1016/j.neuroimage.2013.10.067

Hickey, C., Acunzo, D., & Dell, J. (2023). Suppressive Control of Incentive Salience in Real-World Human Vision. Journal of Neuroscience, 43(37), 6415–6429. 10.1523/JNEUROSCI.0766-23.2023

Hickey, C., Chelazzi, L., & Theeuwes, J. (2014). Reward-Priming of Location in Visual Search. PLoS ONE, 9(7), e103372. 10.1371/journal.pone.0103372

Hickey, C., Di Lollo, V., & McDonald, J. J. (2009). Electrophysiological Indices of Target and Distractor Processing in Visual Search. Journal of Cognitive Neuroscience, 21(4), 760– 775. 10.1162/jocn.2009.21039

Hikosaka, O., Takikawa, Y., & Kawagoe, R. (2000). Role of the Basal Ganglia in the Control of Purposive Saccadic Eye Movements. Physiological Reviews, 80(3), 953–978. 10.1152/physrev.2000.80.3.953

Hillyard, S. A., & Anllo-Vento, L. (1998). Event-related brain potentials in the study of visual selective attention. Proceedings of the National Academy of Sciences, 95(3), 781–787. 10.1073/pnas.95.3.781

Hopfinger, J. B., Woldorff, M. G., Fletcher, E. M., & Mangun, G. R. (2001). Dissociating top-down attentional control from selective perception and action. Neuropsychologia, 39(12), 1277–1291. 10.1016/S0028-3932(01)00117-8

Hyvarinen, A. (1999). Fast and robust fixed-point algorithms for independent component analysis. IEEE Transactions on Neural Networks, 10(3), 626–634. IEEE Transactions on Neural Networks. 10.1109/72.761722

Ipata, A. E., Gee, A. L., Gottlieb, J., Bisley, J. W., & Goldberg, M. E. (2006). LIP responses to a popout stimulus are reduced if it is overtly ignored. Nature Neuroscience, 9(8), 1071– 1076. 10.1038/nn1734

Jiang, Y. V., Swallow, K. M., Rosenbaum, G. M., & Herzig, C. (2013). Rapid acquisition but slow extinction of an attentional bias in space. Journal of Experimental Psychology: Human Perception and Performance, 39(1), 87–99. 10.1037/a0027611

King, J.-R., & Dehaene, S. (2014). Characterizing the dynamics of mental representations: The temporal generalization method. Trends in Cognitive Sciences, 18(4), 203–210. 10.1016/j.tics.2014.01.002

Klink, P. C., Teeuwen, R. R. M., Lorteije, J. A. M., & Roelfsema, P. R. (2023). Inversion of pop-out for a distracting feature dimension in monkey visual cortex. Proceedings of the National Academy of Sciences, 120(9), e2210839120. 10.1073/pnas.2210839120

Krauzlis, R. J., Lovejoy, L. P., & Zénon, A. (2013). Superior Colliculus and Visual Spatial Attention. Annual Review of Neuroscience, 36(1), 165–182. 10.1146/annurev-neuro-062012-170249

Lega, C., Ferrante, O., Marini, F., Santandrea, E., Cattaneo, L., & Chelazzi, L. (2019). Probing the Neural Mechanisms for Distractor Filtering and Their History-Contingent Modulation by Means of TMS. The Journal of Neuroscience, 39(38), 7591–7603. 10.1523/JNEUROSCI.2740-18.2019

Lega, C., Santandrea, E., Ferrante, O., Serpe, R., Dolci, C., Baldini, E., Cattaneo, L., & Chelazzi, L. (2020). Modulating the influence of recent trial history on attentional capture via transcranial magnetic stimulation (TMS) of right TPJ. Cortex, 133, 149–160. 10.1016/j.cortex.2020.09.009

Li, Z. (2002). A saliency map in primary visual cortex. Trends in Cognitive Sciences, 6(1), 9–16.

Liesefeld, H. R., Lamy, D., Gaspelin, N., Geng, J. J., Kerzel, D., Schall, J. D., Allen, H. A., Anderson, B. A., Boettcher, S., Busch, N. A., Carlisle, N. B., Colonius, H., Draschkow, D., Egeth, H., Leber, A. B., Müller, H. J., Röer, J. P., Schubö, A., Slagter, H. A., … Wolfe, J. (2024). Terms of debate: Consensus definitions to guide the scientific discourse on visual distraction. Attention, Perception, & Psychophysics. 10.3758/s13414-023-02820-3

Liesefeld, H. R., & Müller, H. J. (2021). Modulations of saliency signals at two hierarchical levels of priority computation revealed by spatial statistical distractor learning. Journal of Experimental Psychology: General, 150(4), 710–728. 10.1037/xge0000970

Lin, R., Meng, X., Chen, F., Li, X., Jensen, O., Theeuwes, J., & Wang, B. (2024). Neural evidence for attentional capture by salient distractors. Nature Human Behaviour, 8(5), 932–944. 10.1038/s41562-024-01852-5

Lu, R., Dermody, N., Duncan, J., & Woolgar, A. (2024). Aperiodic and oscillatory systems underpinning human domain-general cognition. Communications Biology, 7(1), 1643. 10.1038/s42003-024-07397-7

Luck, S. J., Gaspelin, N., Folk, C. L., Remington, R. W., & Theeuwes, J. (2021). Progress toward resolving the attentional capture debate. Visual Cognition, 29(1), 1–21. 10.1080/13506285.2020.1848949

Luck, S. J., & Hillyard, S. A. (1994). Electrophysiological correlates of feature analysis during visual search. Psychophysiology, 31(3), 291–308. 10.1111/j.1469-8986.1994.tb02218.x

Marini, F., Demeter, E., Roberts, K. C., Chelazzi, L., & Woldorff, M. G. (2016). Orchestrating Proactive and Reactive Mechanisms for Filtering Distracting Information: Brain-Behavior Relationships Revealed by a Mixed-Design fMRI Study. The Journal of Neuroscience, 36(3), 988–1000. 10.1523/JNEUROSCI.2966-15.2016

Maris, E., & Oostenveld, R. (2007). Nonparametric statistical testing of EEG- and MEG-data. Journal of Neuroscience Methods, 164(1), 177–190. 10.1016/j.jneumeth.2007.03.024

Moher, J., & Egeth, H. E. (2012). The ignoring paradox: Cueing distractor features leads first to selection, then to inhibition of to-be-ignored items. Attention, Perception, & Psychophysics, 74(8), 1590–1605. 10.3758/s13414-012-0358-0

Munneke, J., Van Der Stigchel, S., & Theeuwes, J. (2008). Cueing the location of a distractor: An inhibitory mechanism of spatial attention? Acta Psychologica, 129(1), 101–107. 10.1016/j.actpsy.2008.05.004

Niso, G., Gorgolewski, K. J., Bock, E., Brooks, T. L., Flandin, G., Gramfort, A., Henson, R. N., Jas, M., Litvak, V., T. Moreau, J., Oostenveld, R., Schoffelen, J.-M., Tadel, F., Wexler, J., & Baillet, S. (2018). MEG-BIDS, the brain imaging data structure extended to magnetoencephalography. Scientific Data, 5(1), 180110. 10.1038/sdata.2018.110

Pedregosa, F., Varoquaux, G., Gramfort, A., Michel, V., Thirion, B., Grisel, O., Blondel, M., Prettenhofer, P., Weiss, R., Dubourg, V., Vanderplas, J., Passos, A., Cournapeau, D., Brucher, M., Perrot, M., & Duchesnay, É. (2011). Scikit-learn: Machine Learning in Python. Journal of Machine Learning Research, 12(85), 2825–2830.

Perruchet, P., & Pacton, S. (2006). Implicit learning and statistical learning: One phenomenon, two approaches. Trends in Cognitive Sciences, 10(5), 233–238. 10.1016/j.tics.2006.03.006

Pouget, A., Beck, J. M., Ma, W. J., & Latham, P. E. (2013). Probabilistic brains: Knowns and unknowns. Nature Neuroscience, 16(9), 1170–1178. 10.1038/nn.3495

Reeder, R. R., Olivers, C. N. L., & Pollmann, S. (2017). Cortical evidence for negative search templates. Visual Cognition, 25(1–3), 278–290. 10.1080/13506285.2017.1339755

Richter, D., van Moorselaar, D., & Theeuwes, J. (2024). Proactive distractor suppression in early visual cortex. bioRxiv.

Ruff, C. C., & Driver, J. (2006). Attentional Preparation for a Lateralized Visual Distractor: Behavioral and fMRI Evidence. Journal of Cognitive Neuroscience, 18(4), 522–538. 10.1162/jocn.2006.18.4.522

Sauter, M., Liesefeld, H. R., Zehetleitner, M., & Müller, H. J. (2018). Region-based shielding of visual search from salient distractors: Target detection is impaired with same- but not different-dimension distractors. Attention, Perception, & Psychophysics, 80(3), 622– 642. 10.3758/s13414-017-1477-4

Seidl, K. N., Peelen, M. V., & Kastner, S. (2012). Neural Evidence for Distracter Suppression during Visual Search in Real-World Scenes. The Journal of Neuroscience, 32(34), 11812– 11819. 10.1523/JNEUROSCI.1693-12.2012

Seijdel, N., Marshall, T. R., & Drijvers, L. (2023). Rapid invisible frequency tagging (RIFT): A promising technique to study neural and cognitive processing using naturalistic paradigms. Cerebral Cortex, 33(5), 1626–1629. 10.1093/cercor/bhac160

Sherman, B. E., Graves, K. N., & Turk-Browne, N. B. (2020). The prevalence and importance of statistical learning in human cognition and behavior. Current Opinion in Behavioral Sciences, 32, 15–20. 10.1016/j.cobeha.2020.01.015

Sprague, T. C., & Serences, J. T. (2013). Attention modulates spatial priority maps in the human occipital, parietal and frontal cortices. Nature Neuroscience, 16(12), 1879–1887. 10.1038/nn.3574

Stokes, M. G. (2015). ‘Activity-silent’ working memory in prefrontal cortex: A dynamic coding framework. Trends in Cognitive Sciences, 19(7), 394–405. 10.1016/j.tics.2015.05.004

Stokes, M. G., Kusunoki, M., Sigala, N., Nili, H., Gaffan, D., & Duncan, J. (2013). Dynamic Coding for Cognitive Control in Prefrontal Cortex. Neuron, 78(2), 364–375. 10.1016/j.neuron.2013.01.039

Suzuki, M., & Gottlieb, J. (2013). Distinct neural mechanisms of distractor suppression in the frontal and parietal lobe. Nature Neuroscience, 16(1), 98–104. 10.1038/nn.3282

Taulu, S., Kajola, M., & Simola, J. (2003). Suppression of Interference and Artifacts by the Signal Space Separation Method. Brain Topography, 16(4), 269–275. 10.1023/B:BRAT.0000032864.93890.f9

Theeuwes, J. (1992). Perceptual selectivity for color and form. Perception & Psychophysics, 51(6), 599–606. 10.3758/BF03211656

Theeuwes, J. (2010). Top–down and bottom–up control of visual selection. Acta Psychologica, 135(2), 77–99. 10.1016/j.actpsy.2010.02.006

Theeuwes, J., Bogaerts, L., & Van Moorselaar, D. (2022). What to expect where and when: How statistical learning drives visual selection. Trends in Cognitive Sciences, 26(10), 860– 872. 10.1016/j.tics.2022.06.001

Turatto, M., & Pascucci, D. (2016). Short-term and long-term plasticity in the visual-attention system: Evidence from habituation of attentional capture. Neurobiology of Learning and Memory, 130, 159–169. 10.1016/j.nlm.2016.02.010

Van Moorselaar, D., & Slagter, H. A. (2019). Learning What Is Irrelevant or Relevant: Expectations Facilitate Distractor Inhibition and Target Facilitation through Distinct Neural Mechanisms. The Journal of Neuroscience, 39(35), 6953–6967. 10.1523/JNEUROSCI.0593-19.2019

Van Moorselaar, D., & Slagter, H. A. (2020). Inhibition in selective attention. Annals of the New York Academy of Sciences, 1464(1), 204–221. 10.1111/nyas.14304

Wang, B., & Theeuwes, J. (2018a). How to inhibit a distractor location? Statistical learning versus active, top-down suppression. Attention, Perception, & Psychophysics, 80(4), 860–870. 10.3758/s13414-018-1493-z

Wang, B., & Theeuwes, J. (2018b). Statistical regularities modulate attentional capture. Journal of Experimental Psychology: Human Perception and Performance, 44(1), 13.

Wang, B., Van Driel, J., Ort, E., & Theeuwes, J. (2019). Anticipatory Distractor Suppression Elicited by Statistical Regularities in Visual Search. Journal of Cognitive Neuroscience, 31(10), 1535–1548. 10.1162/jocn_a_01433

Wang, J.-Z., Williamson, S. J., & Kaufman, L. (1992). Magnetic source images determined by a lead-field analysis: The unique minimum-norm least-squares estimation. IEEE Transactions on Biomedical Engineering, 39(7), 665–675. 10.1109/10.142641

Zhang, B., Weidner, R., Allenmark, F., Bertleff, S., Fink, G. R., Shi, Z., & Müller, H. J. (2022). Statistical Learning of Frequent Distractor Locations in Visual Search Involves Regional Signal Suppression in Early Visual Cortex. Cerebral Cortex, 32(13), 2729–2744. 10.1093/cercor/bhab377

Zhigalov, A., Herring, J. D., Herpers, J., Bergmann, T. O., & Jensen, O. (2019). Probing cortical excitability using rapid frequency tagging. NeuroImage, 195, 59–66. 10.1016/j.neuroimage.2019.03.056

